# Aberrant DNA methylation is co-regulated across the genome in leukemia and other types of cancer

**DOI:** 10.64898/2026.06.05.729844

**Authors:** Monica Varona Baranda, Sven Liesenfelder, Florian Kraft, Chao-Chung Kuo, Juan-Felipe Perez-Correa, Edgar Jost, Thomas Stiehl, Wolfgang Wagner

## Abstract

Epigenetic dysregulation is a defining feature of cancer, but it remains poorly understood how this is coordinated across the genome. In this study we focusd on DNA methylation (DNAm) in acute myeloid leukemia (AML). Despite highly heterogeneous and largely patient-specific patterns, we identified co-regulated clusters of CpGs that could be assembled into reproducible epigenetic networks. Multilinear regression models accurately predicted the patient-specific DNAm deviations, even for CpGs located on different chromosomes. The alterations were mirrored on homologous chromosomes and there was no clear association with epigenetic driver mutations. Furthermore, we found very similar co-regulation patterns in acute lymphoblastic leukemia (ALL); with AML-derived models successfully predicting the ALL-associated DNAm changes. Notably, the top 1000 AML-associated CpGs showed also pronounced aberrations in DNAm levels across 46 other cancer types, whereas this was hardly observed across multiple non-malignant cell types. Co-regulation analysis realed very similar patterns in non-malignant blood and pan-cancer analysis, albeit the DNAm levels remained overall consistent in the controls. Collectively, our findings demonstrate that the complex, patient-specific DNAm landscapes observed in leukemia are not random. Instead, they are orchestrated within expanded epigenetic networks, which also exist in non-maligant cells, highlighting a higher-order regulatory layer in cancer epigenomics.

## Introduction

Cancer is not only associated with genetic mutations but also with extensive epigenetic re-programming (1). Among the epigenetic mechanisms, DNA methylation (DNAm) at CpG dinucleotides is especially well studied because it is relatively stable, amenable to high-throughput profiling, and generally disturbed in cancer (2). Whole-genome and targeted methylome analyses repeatedly demonstrate that the DNAm profile is dramatically reshaped in virtually every malignant disease (3). These alterations can be harnessed to create epigenetic signatures that serve as diagnostic, prognostic, or predictive biomarkers (4, 5). In several pan-cancer studies, a subset of epigenetic aberrations has even been reported to recur across biologically diverse tumour types (6, 7).

The molecular mechanisms that shape cancer-associated DNAm patterns are not yet fully understood. Most investigations compare cancerous *versus* non-cancerous tissue to identify disease-related DNAm marks. However, many of the detected differences may simply arise from shifts in cellular composition relative to healthy tissue (8-10). Moreover, because tumours are clonally derived, they initially capture the epigenetic makeup of the tumor-initiating cell. Therefore, some CpGs initially reflect extreme DNAm values that may drift toward hemi-methylated states due to stochastic changes as the disease progresses (11). Patient-specific DNAm patterns can also be driven by somatic mutations in epigenetic regulators such as DNMT3A or TET2, although this only applies to some of cases (12, 13). Furthermore, mutations may generally impact DNAm in their immediate genomic vicinity (14, 15). Other studies propose that common transcription-factor binding sites, chromatin-state domains, or three-dimensional DNA contacts shape the patient-specific methylation landscape (16). However, so far it is unclear if epigenetic aberrations arise by local mechanisms, or if they are rather concerted across the genome in coordinated epigenetic networks with co-regulated hubs (16, 17).

In the present work we addressed this question in acute myeloid leukemia (AML), a disease characterised by a high proportion of malignant cells (as estimated by blast counts) and for which DNAm profiles of non-malignant leukocyte subsets are readily available. AML is a heterogeneous entity, and numerous DNAm signatures have been proposed to discriminate favourable from adverse risk groups (18-20). Aberrant DNAm profiles appear to be patient-specific, and we have previously employed bisulphite amplicon sequencing of regions with abnormal methylation to detect patient-specific patterns that could serve as measurable residual disease (MRD) markers (21). Here, we compiled DNAm datasets of AML patients and healthy controls to investigate how AML-associated methylation changes are coordinated across the genome. We demonstrate that patient-specific DNAm aberrations are not random but are organised into reproducible epigenetic networks, which show also pronounced aberrations in many other types of cancer, and reveal similar co-regulation even in non-malignant blood samples. This finding challenges the prevailing view of methylation changes as isolated events and supports a model in which they function as components of a coordinated epigenetic network.

## Results

### AML-associated DNA methylation changes in co-regulation networks

To select AML-associated CpGs, we initially compiled an identification set of 344 DNAm profiles of AML patients (five studies) alongside 1693 healthy blood samples for control (3 studies, all 450K, Supplemental Table S1). We focused on CpGs, which are either highly methylated (mean DNAm > 90%) or non-methylated (mean DNAm <10%) in the controls, to provide a defined ground state of DNAm that is less susceptible to reflect the epigenetic makeup of the tumor-initiating cell (11). Subsequently, we ranked these CpGs by their absolute mean DNAm difference in AML *versus* controls and excluded CpGs in close proximity (<5 kb to each other), since neighboring CpGs may be simultaneously changed by local means within differentially methylated regions (Figure 1A). Based on this, we identified the top 1000 CpGs, where 68 CpGs lost and 932 CpGs gained aberrant DNAm in the AML cohort (Figure 1B; Supplemental Table S2). When we used an independent validation set of 393 AML samples and 1237 controls (3 and 2 studies, respectively, all EPIC, Supplemental Table S1), we observed overall consistent changes (Figure 1C). Notably, these AML-associated aberrations exhibit high variability between patients (Figure 1D). In contrast, none of these CpGs showed pronounced differences in DNAm between purified leukocyte subsets (Supplemental Figure S1A). Thus, we identified AML-associated CpGs that are either methylated or non-methylated in controls, but highly variable between AML patient samples.

**Figure 1:**
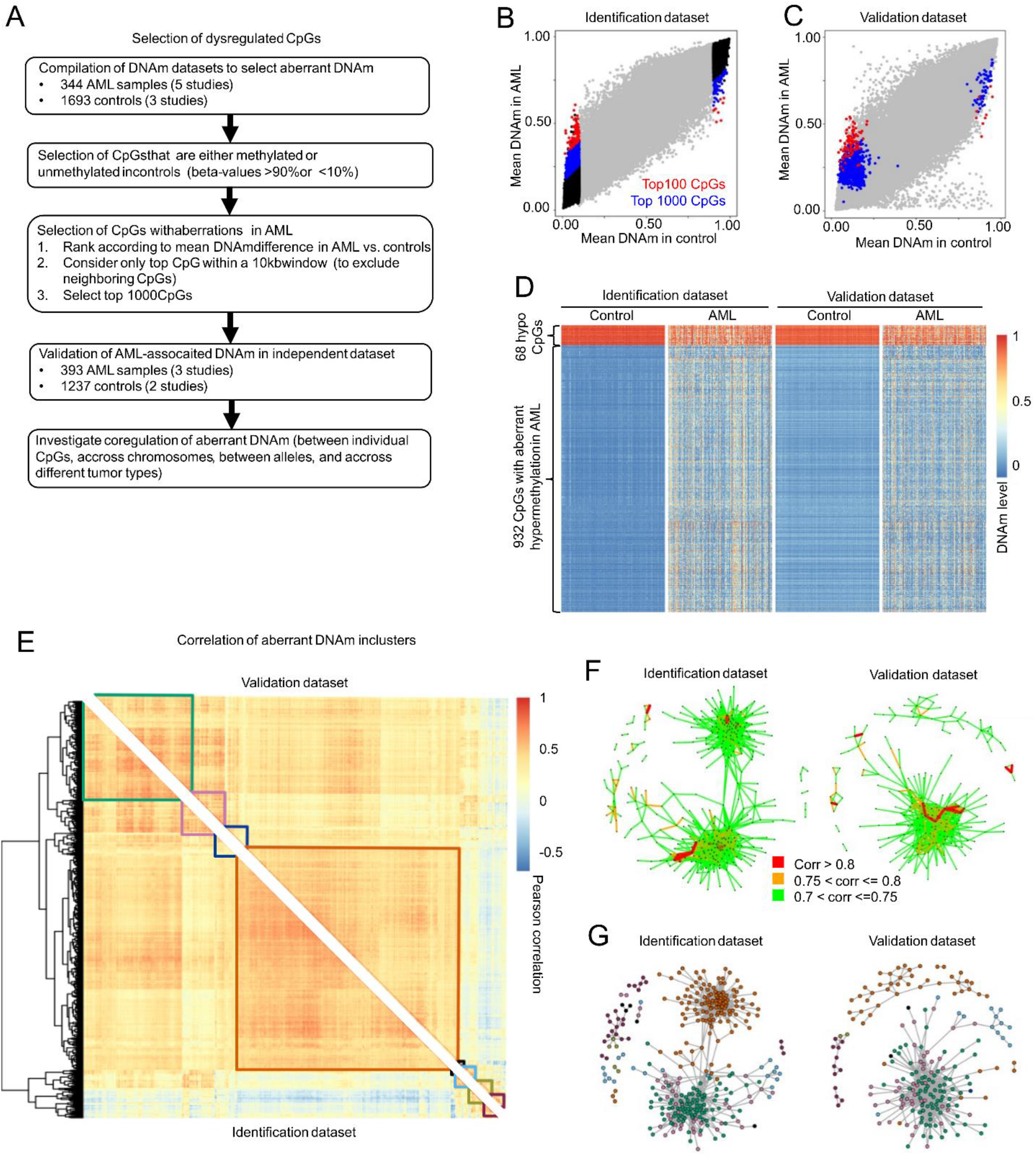
Dysregulated DNAm in AML is co-regulated in clusters. **A)** Selection process of aberrant DNAm in AML. **B**,**C)** Scatterplots of the mean DNAm (beta-value) differences between healthy controls and AML in (B) the identification datasets, and in (C) the validation dataset. The selected top 100 (red), top 1000 CpGs (blue), and stable CpGs in healthy samples (black) are depicted. **D)** Heatmaps demonstrating heterogeneity of DNAm levels at the top 1000 AML-associated CpGs (including 68 CpGs with aberrant hypo- and 932 with aberrant hypermethylation in AML). For graphical presentation 300 randomly chosen profiles were depicted from controls and AML patients in both datasets. **E)** Correlation analysis (Pearson’s correlation) of DNAm levels between each of the top 1000 AML-associated CpGs across the individual patients of the identification dataset and the validation dataset. Hierarchical clustering of this correlation matrix is based on the identification dataset, and eight clusters of correlating CpGs are highlighted. **F)** Network plots for identification (left) and validation (right) dataset. Each dot represents one of the top 1000 CpG and only pairs with a correlation value higher than 0.7 are displayed. **G)** The same network plots where nodes are colored according to the clusters identified in (E).

Principal component analysis (PCA) of the top 1000 AML-associated CpGs did not cluster patients according to study, microarray platform, French-American-British (FAB) classification of AML, AML subtypes, or risk stratification (Supplemental Figure S1B-F). The CpGs with aberrant gain and loss in AML were equally distributed across all chromosomes (Supplemental Figure S2G). They were moderately enriched in CpG-islands (CGI), in promotor regions, and the first exons (Supplemental Figure S2H-I). Furthermore, these 1000 CpGs were enriched in genes related to functional categories in neuronal differentiation, which might possibly reflect to mesenchymal-to-epithelial transition (Supplemental Figure S2J) (22).

To explore if the heterogeneous AML-associated DNAm might be co-regulated between CpGs, we initially performed pair-wise correlation analysis at individual CpGs within AML samples (Pearson’s correlation). While many pairs of CpGs did not reveal high correlation, we identified eight clusters of highly correlated CpGs. Notably, almost the exact same correlation pattern was observed in the independent validation set, indicating that this co-regulation is highly preserved in AML (Figure 1E). Next we performed network visualization of the top 1000 CpGs, where each node corresponds to a CpG with significant pairwise correlations of R > 0.7 (Figure 1F). The co-regulated subnetworks corresponded to the clusters identified by the correlation matrix (Figure 1G). This analysis highlighted that aberrant DNAm is acquired genome wide in a very heterogeneous and patient-specific manner, but the DNAm patterns seem to be concerted in hubs of co-ocurrence across independent AML cohorts.

### Predicting AML-associated DNAm patterns on independent chromosomes

To further substantiate the hypothesis that AML-associated DNAm patterns are co-regulated in a patient-specific manner, we trained multilinear models to predict DNAm levels of CpGs on a given chromosome based on the AML-associated CpGs on the other chromosomes. To this end, one third of the identification dataset (Supplemental Table S1) was used to train elastic net-based models for each CpGs based on the top 1000 AML-associated CpGs remaining on other chromosomes (Figure 2A). The predictions showed overall high correlation with the observed DNAm levels in the training set, in the remaining two thirds that was used as *testing* set, and in the previous validation set of 393 independent AML samples (Figure 2B,C). The DNAm patterns at the AML-associated CpGs reveal patient-specific patterns, which can be predicted based on the DNAm profiles on other chromosomes of the same patient (Figure 2D,E). This analysis demonstrated that the AML-associated DNAm patterns do not arise independently in a stochastic manner, but that they are concerted across different chromosomes.

**Figure 2:**
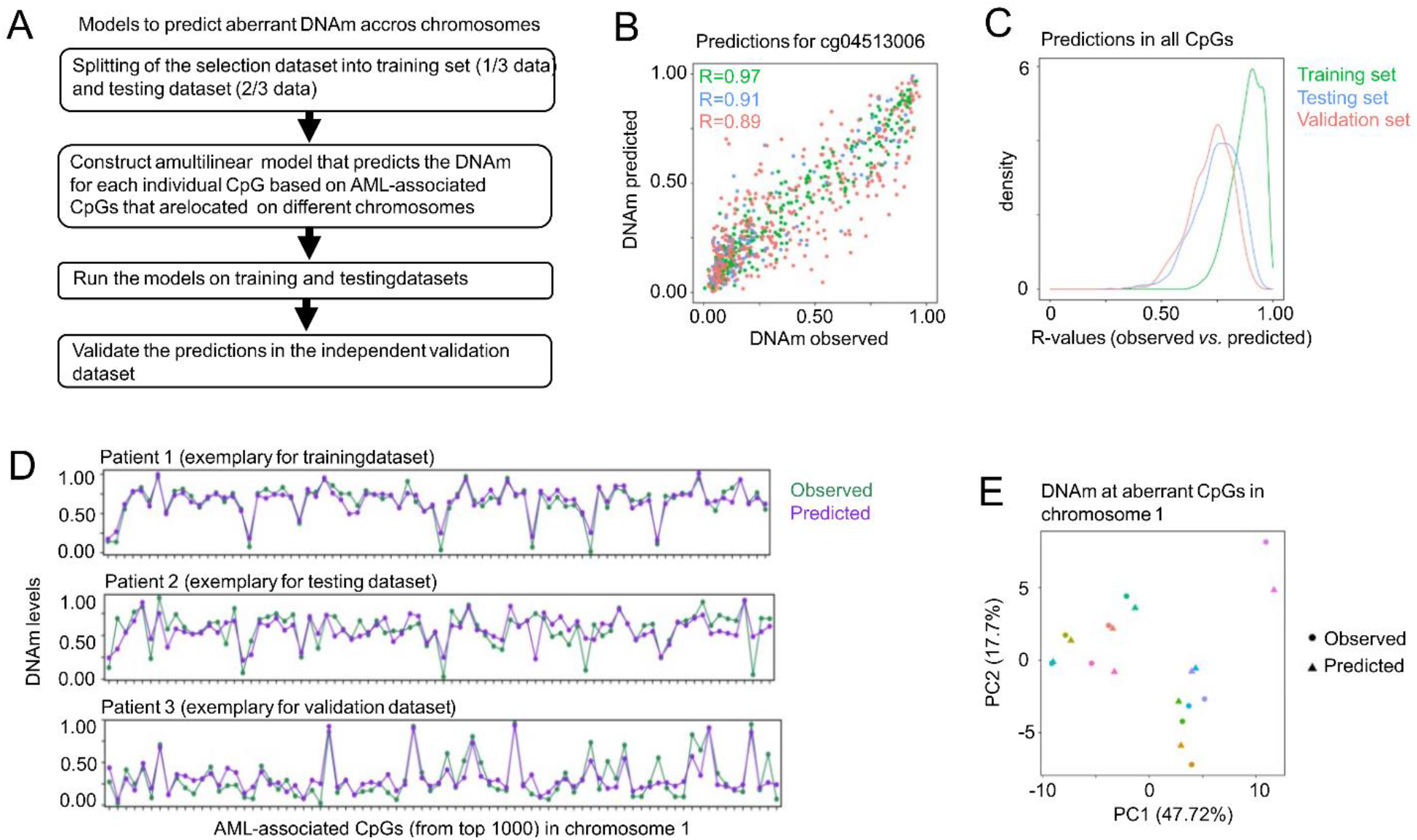
Dysregulated DNA methylation patterns can be predicted across chromosomes. **A)** Schematic representation on how multilinear models were trained within the top 1000 AML-associated CpGs to predict aberrant DNAm levels on independent chromosomes. **B)** Exemplary scatterplot for the CpG cg04513006 (on chromosome 16) to demonstrate observed DNAm levels and the predicted DNAm levels based on a elastic net-regularized multilinear model of CpGs in the remaining chromosomes (chr. 1 to 15, and 17 to 22). Pearson’s correlation (R) is provided within the training (green), testing (blue), and independent validation dataset (orange). **C)** Density plot of the correlation values between observed and predicted DNAm values for all CpGs in the different datasets. **D)** To demonstrate the heterogeneity of dysregulated DNAm between AML patients, and to visualize that these patterns can be reproduced based on multilinear models using aberrant CpGs on independent chromosomes, the observed and predicted DNAm levels are exemplarily depicted for the 80 CpGs on chromosome 1 (within the top 1000 CpGs) for three exemplary AML patients. **E)** Principal component analysis for 10 randomly selected AML samples (indicated by different colors) to demonstrate that predicted (triangle) and observed (circle) DNAm patterns at these 80 CpGs at chromosome 1 are clustering together in a patient-specific manner.

### AML-associated DNAm is assimilated on both alleles

To gain insight if the AML-associated CpGs are located within differentially methylated regions (DMRs) and if aberrations affect both alleles, we used nanopore sequencing data of 10 AML and 9 control samples with adaptive sampling sequencing to enrich the 100 top AML-associated CpGs (Supplemental Figure 2A). This analysis demonstrated that aberrant DNAm is often in DMRs of about 1kb (Figure 3A,B).

**Figure 3:**
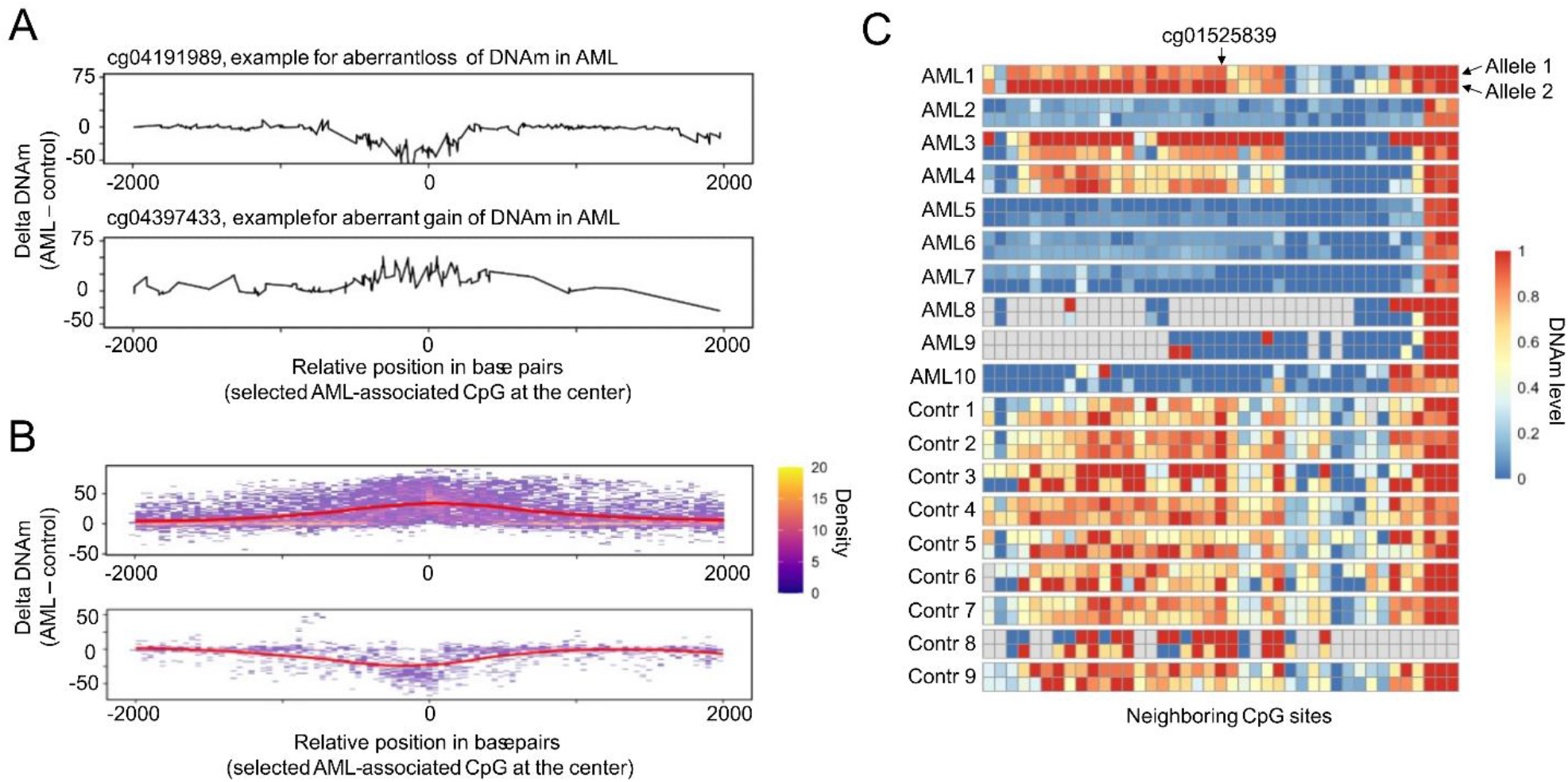
Dysregulated DNAm patterns affect both alleles. **A**,**B)** To determine if the dysregulated CpGs fall into differentially methylated regions (DMRs), we used adaptive nanopore sequencing around the top 100 AML-associated CpGs. (A) Exemplary line plots depict the difference of mean DNAm levels between 10 AML and 9 control samples for two CpGs (of the top 1000 AML-associated CpGs) and their neighboring CpGs within a 4kb window. (B) Overlay of such analysis at the top 100 AML-associated CpGs demonstrates that dysregulated CpGs are typically in DMRs ranging about 1000 bp. **C)** To compare the DNAm patterns on both alleles, we used phased nanopore sequencing data and the patterns are exemplarily depicted for cg01525839 and all neighboring CpGs within a 4kb window (grey boxes indicate CpGs without coverage in the corresponding sample).

Additionally, phased nanopore sequencing data was used to assign the reads to the same homologous chromosome, to understand whether aberrant DNAm dynamics on different haplotypes are independent (11), or if they are assimilated (21). We observed that the AML-associated DNAm patterns are overall very similar on the homologous chromosomes (Figure 3C, Supplemental Figure 2B,C), indicating that aberrant DNAm patterns are either assimilated between the haplotypes or coordinated in the same way across both alleles.

### AML-associated DNAm has moderate association with clinical outcome

To explore if the overall changes at our AML-associated CpGs have prognostic relevance, we focused on the AML dataset of the cancer genome atlas (TCGA) (23). The top 1000 CpGs were grouped into the 68 CpGs with aberrant hypomethylation and the 932 CpGs with aberrant hypermethylation to determine the median DNAm value at these sites for each sample (Figure 4A,B; Supplemental Figure 3A,B). Notably, the blast counts were only moderately associated with the median DNAm changes (R = 0.3 for CpGs with aberrant hypermethylation; Pearson’s correlation), indicating that our signature does not simply reflect tumor burden (Figure 4C,D). About half of the AML samples have normal karyotype, often without structural abnormalities (23), and we observed that particularly the median AML-associated hypomethylation was more pronounced in samples with a lower fraction of the genome altered (R = 0.3; Figure 4E,F). Genomic mutations in epigenetic writers, such as *ASLX1, RUNX1, DNMT3A, IDH1, IDH2*, and *TET2* did now show significant association with AML-associated DNAm changes at our top 1000 CpGs (Figure 4G,H). We have also analyzed if the aberrant DNAm levels are related to event free survival (Figure 4I,J), or overall survival (Supplemental Figure 3C,D), but we did not observe a significant association – in tendency, more epigenetic aberrations were associated with better outcome after treatment. In a univariable Cox proportional hazards model, higher median AML-associated hypermethylation was significantly associated with longer disease-free survival (HR = 1.22, 95% CI 1.01–1.47, *p* = 0.037), but this association did not remain significant after adjusting for age and gender.

**Figure 4:**
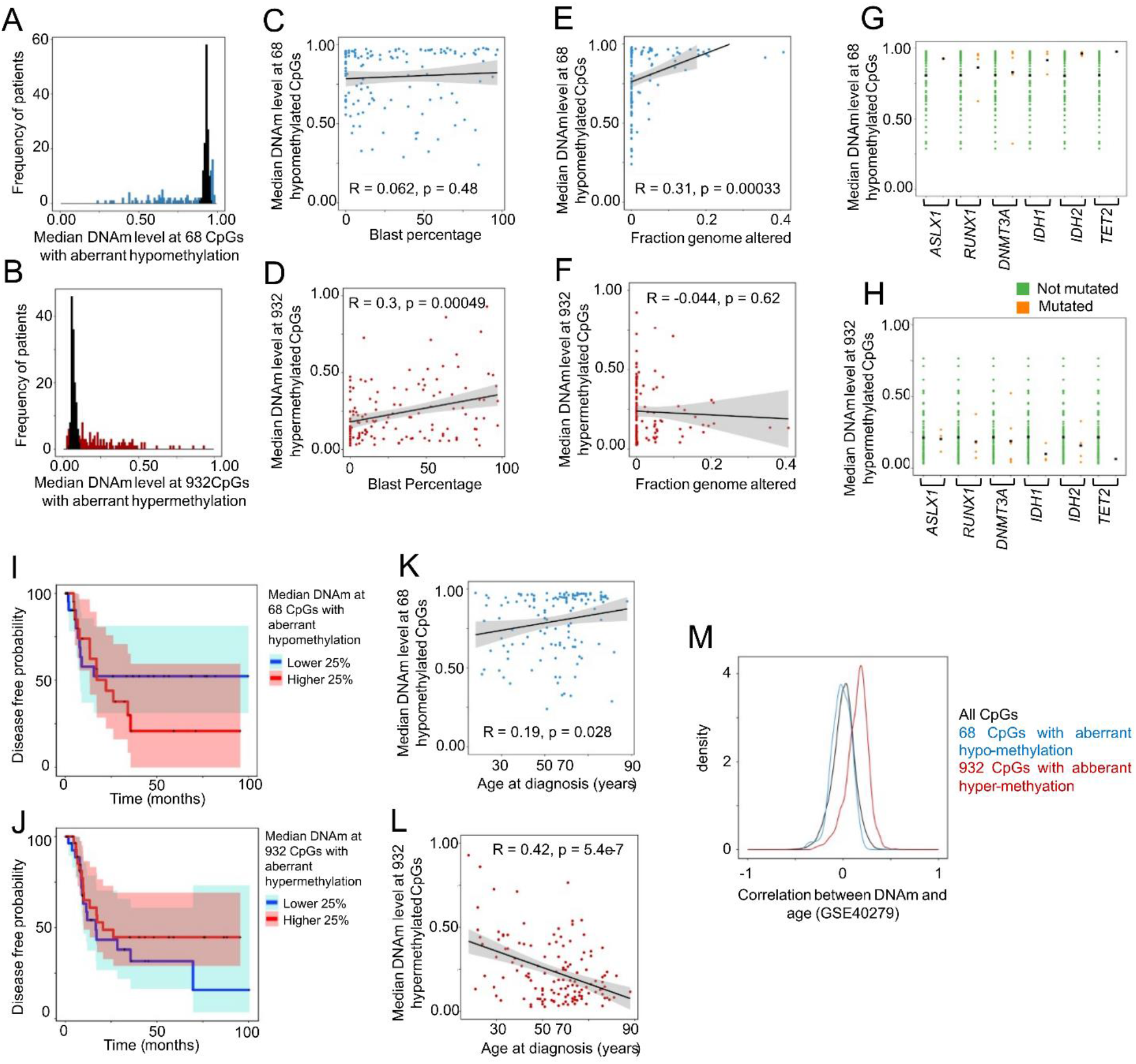
Association of dysregulated DNAm with patient characteristics. **A**,**B)** Histogram with the frequency of median DNAm levels within the (A) 68 CpGs with aberrant hypomethylation in AML (blue), and (B) 932 CpGs with aberrant hypermethylation in AML (red). The median DNAm levels are provided for AML patients of the TCGA cohort (23), and for comparison for healthy controls (black). **C**,**D)** Correlation between median DNAm at CpGs with aberrant hypo- (C), and hypermethylation in AML (D) in comparison to blast counts in peripheral blood. There was a moderate association with blast counts at the hypermethylated CpGs (p-values determined by stat_cor function from package ggpubr). **E**,**F)** Correlation between median DNAm levels at CpGs with aberrant hypo- (E) and hyper-methylation (F) in comparison to the fraction of the genome altered, as provided by the TCGA, indicating that AML samples with more genomic aberrations have less DNAm dysregulation at CpGs with aberrant hypomethylation. **G**,**H)** To explore if the dysregulation at the top 1000 AML-associated CpGs could be attributed to mutations in epigenetic writers, we compared the median DNAm at the CpGs with aberrant loss (G) or gain in DNAm (H) in samples with or without mutations in six epigenetic writers. Notably, in samples without such mutations, median DNAm per patient for aberrant hyper- and hypomethylated CpGs in AML appeared in tendency even higher (median of all patients in each category indicated by black dot). **I**,**J)** Kaplan-Meier plots showing the disease-free probability in the quartile with the lowest (blue) or highest (red) median DNAm at either the CpGs with aberrant loss of DNAm (I) or aberrant gain of DNAm in AML (J). In tendency, both curves indicate that samples with more epigenetic dysregulation have a better prognosis, but the analysis did not reach statistical significance. **K**,**L)** Correlation between median DNAm levels at CpGs with aberrant loss (K) or gain of DNAm (L) in comparison to the age at diagnosis. Younger patients reveal more median dysregulation at these CpGs. **M)** To investigate if CpGs with aberrant loss (blue) or gain of DNAm in AML (red) are enriched at age-associated CpGs, we used DNAm profiles from a dataset of healthy blood samples (GSE40279) to identify the correlation with chronological age for each CpGs. The density plot demonstrates that particularly the CpGs with aberrant gain in methylation are also gaining DNAm during aging as compared to all CpGs on the array that passed quality control (p-value < 10^−15^; Wilcox test).

Notably, younger AML patients showed, in tendency, higher epigenetic aberrations at hyper- and hypomethylated sites (Figure 4K,L). Next, we investigated if the AML-associated CpGs reveal also age-associated changes in healthy blood – despite the fact that our selection was restricted to sites that are overall non- or highly methylated in controls. When we investigated the correlation of DNAm in a cohort of healthy donors of different ages (24), we observed that particularly the 932 CpGs with aberrant hyper-methylation in AML show also gain of methylation during aging (Figure 4M). Taken together, the median DNAm levels at our AML-associated CpGs do not appear to be suitable as a prognostic biomarker, which might be attributed to the fact, that they occurred rather independent from tumor burden, and that they were particularly high in younger patients without genomic alterations.

### ALL recapitulates co-regulation of DNAm at the AML-associated CpGs

To investigate whether the epigenetic co-regulation network we uncovered in AML also operates in other neoplasms, we exemplarily turned to DNAm datasets of acute lymphoblastic leukemia (ALL; N =152 samples, GSE182313 and GSE147667) (25, 26). Using the same pipeline employed for AML, we extracted ALL-specific DNAm aberrations. Remarkably, all top 1000 CpGs exhibited gain of methylation in ALL (Figure 5A), and most of these sites showed also aberrant hypermethylation in the AML cohort (Figure 5B). Within the top 1000 AML- and ALL-associated CpGs 238 CpGs were significantly overlapping (p < 10^−20^; hypergeometric test; Figure 5C), and when we ranked the CpGs by their mean DNAm difference *versus* controls, we observed concordant ranking across both disease entities (Spearman correlation = 0.26 across all CpGs with p-value < 10^−15^; Figure 5D). Furthermore, the top 1000 AML-associated CpGs revealed a similar co-regulation pattern with approximately preserved clusters also in the ALL dataset (Supplemental Figure 4).

**Figure 5:**
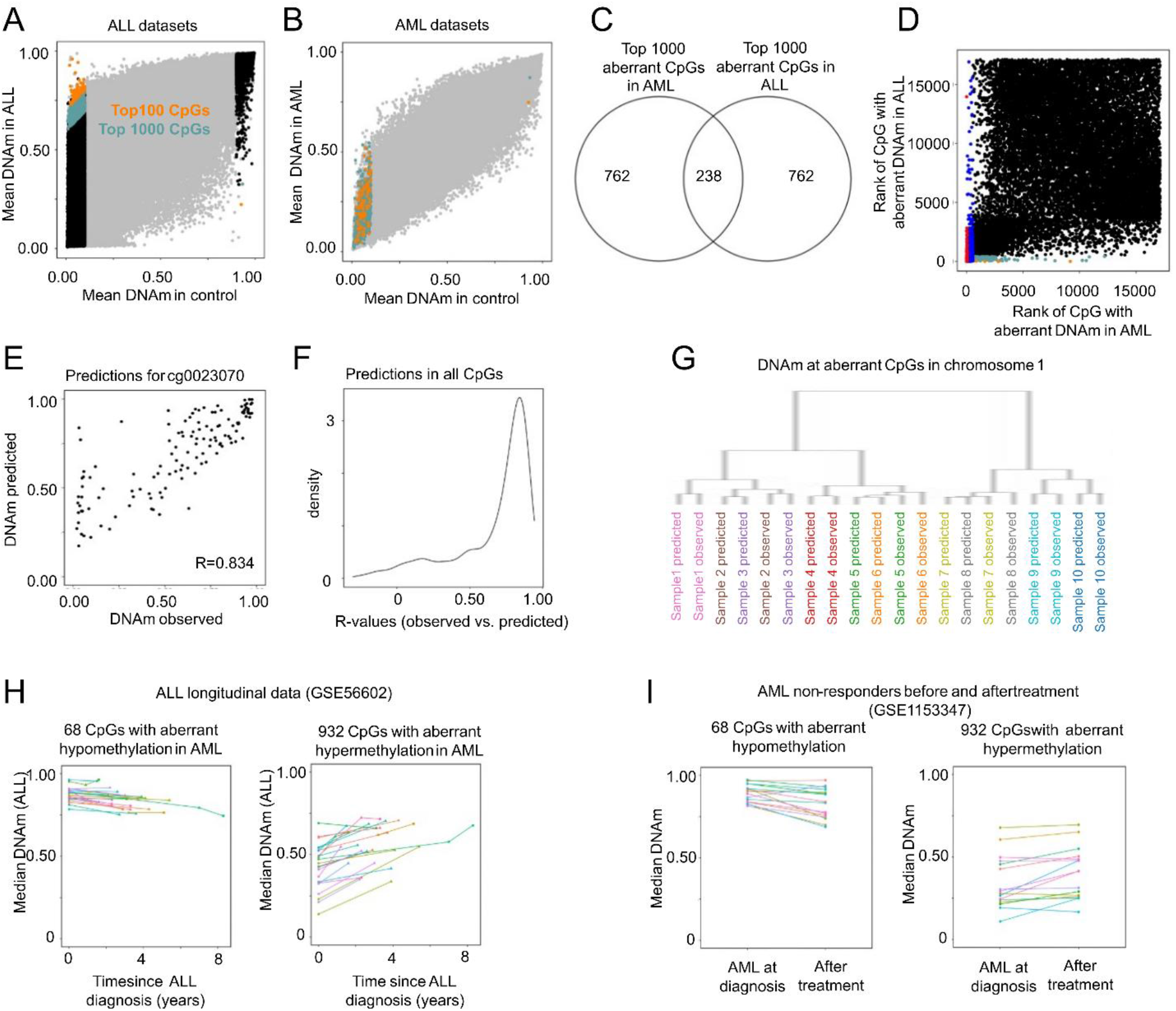
Aberrant DNAm in ALL shows similar co-regulation as in AML. **A)** DNAm datasets of patients with acute lymphoid leukemia (GSE182313 and GSE147667) were used to select CpGs with aberrant DNAm (in analogy to Figure 1A). **B)** Scatterplot displaying the top 100 CpGs (orange) and top 1000 CpGs (green) with aberrant DNAm in ALL in the AML dataset. **C)** Venn Diagram displaying a significant overlap between the top 1000 AML-associated CpGs and the top 1000 ALL-associated CpGs (p < 10^−20^; hypergeometric distribution). **D)** To further compare dysregulation in AML and ALL, we compared their ranks during the selection procedures, with red and blue representing the top100/1000 AML-associated CpGs, and orange and green presenting the top100/1000 ALL-associated CpGs. Overall, a similar set of CpGs showed the highest aberrations in both types of leukemia. **E)** The multilinear prediction model for the CpG site cg0023070, which was trained on AML data and AML-associated CpGs in other chromosomes (as described in Figure 2A,B), was exemplarily used to predict DNAm levels at this site in the ALL datasets. The scatterplot exemplarily demonstrates the correlation of predicted and observed values (R = 0.834, Pearson’s correlation). **F)** Density plot shows the correlation values across all CpGs in the ALL dataset – again using the same prediction models that were previously trained for AML. **G)** Dendogram to demonstrate the relationship of DNAm patterns (observed *versus* predicted) in 80 CpGs on chromosome 1 for 10 randomly selected ALL samples. Albeit the predictors were trained on AML data, the observed and predicted patterns in ALL clustered also relatively close together. **H)** Lineplots displaying the longitudinal changes over different time points in ALL patients (GSE56602) for the median DNAm at CpGs with aberrant loss (left) or gain (right) of DNAm in AML. Overall, the aberrations increased significantly over the years, particularly at CpGs with aberrant hypermethylation. **I)** To estimate if similar longitudinal changes were also observed in AML, we compared the median DNAm in non-responders before and after treatment (GSE1153347), and the aberrations overall increased significantly in the curse of disease (paired t-test: p = 0.0012 for aberrant hypo- and p = 0.0006 for hypermethylated CpGs).

To further explore if the patient-specific co-regulation is preserved between AML and ALL cohorts, we applied, on ALL data, the same elastic-net regression models to predict DNAm on independent chromosomes, which were previously trained on AML data. The same models yielded clear linear relationships between predicted and observed DNAm values for the majority of CpGs also in ALL (788 out of 1000 models have a correlation between observed and predicted values higher than 0.5; Figure 5E,F). Hierarchical clustering of ALL-DNAm profiles based on the AML-associated CpGs on chromosome 1 demonstrated that predicted and observed DNAm profiles clustered together patient-specific, further substanciating that the co-regulatory network architecture identified in AML is overall recapitulated in ALL (Figure 5G).

To determine the dynamics of aberrant DNAm during disease progression, we used longitudinal methylation data from 26 ALL patients (up to 3 time points per patient, GSE56602)(27). This analysis revealed a steady increase in the magnitude of the aberrant methylation signature over the course of disease (Figure 5H). Although comparable longitudinal data was not available for AML, an analysis of diagnosis *versus* post-treatment methylation in non-responders (GSE153347)(28), showed a similar trend of progressive aberrant-methylation with time (Figure 5I, Supplemental Figure 5). Collectively, these findings demonstrate that aberrant DNAm in AML and ALL is coordinated in a similar way and becomes more pronounced as the disease advances.

### Similar epigenetic aberrations are observed in pan-cancer analysis

To investigate whether the DNAm alterations identified in AML reflect a broader oncogenic program, we assembled a pan-cancer cohort comprising 13 additional hematopoietic malignancies and 33 non-hematopoietic tumor types derived from TCGA. For comparison, we additionally compiled a reference panel of 13 non-malignant cell types (Supplemental Table S3). Across the non-malignant reference panel, AML-associated CpG sites displayed largely uniform methylation levels that closely resembled those observed in healthy blood and leukocyte subsets. In contrast, both hematopoietic and solid tumors exhibited widespread aberrant methylation at these CpGs, most prominently at loci that gain DNAm in AML (Figure 6A). Notably, several cancer entities showed in average across their patients even higher median DNAm levels at AML-associated hypermethylated CpGs than those observed in the AML cohort itself (Figure 6B). Moreover, there was a general tendency for cancer entities with poorer 5-year overall survival to exhibit more pronounced DNAm aberrations (Supplemental Figure 6A). Thus, the pronounced and individual AML-associated aberrations are not AML-specific, but these CpGs show even more pronounced aberrations across many other types of cancer.

**Figure 6:**
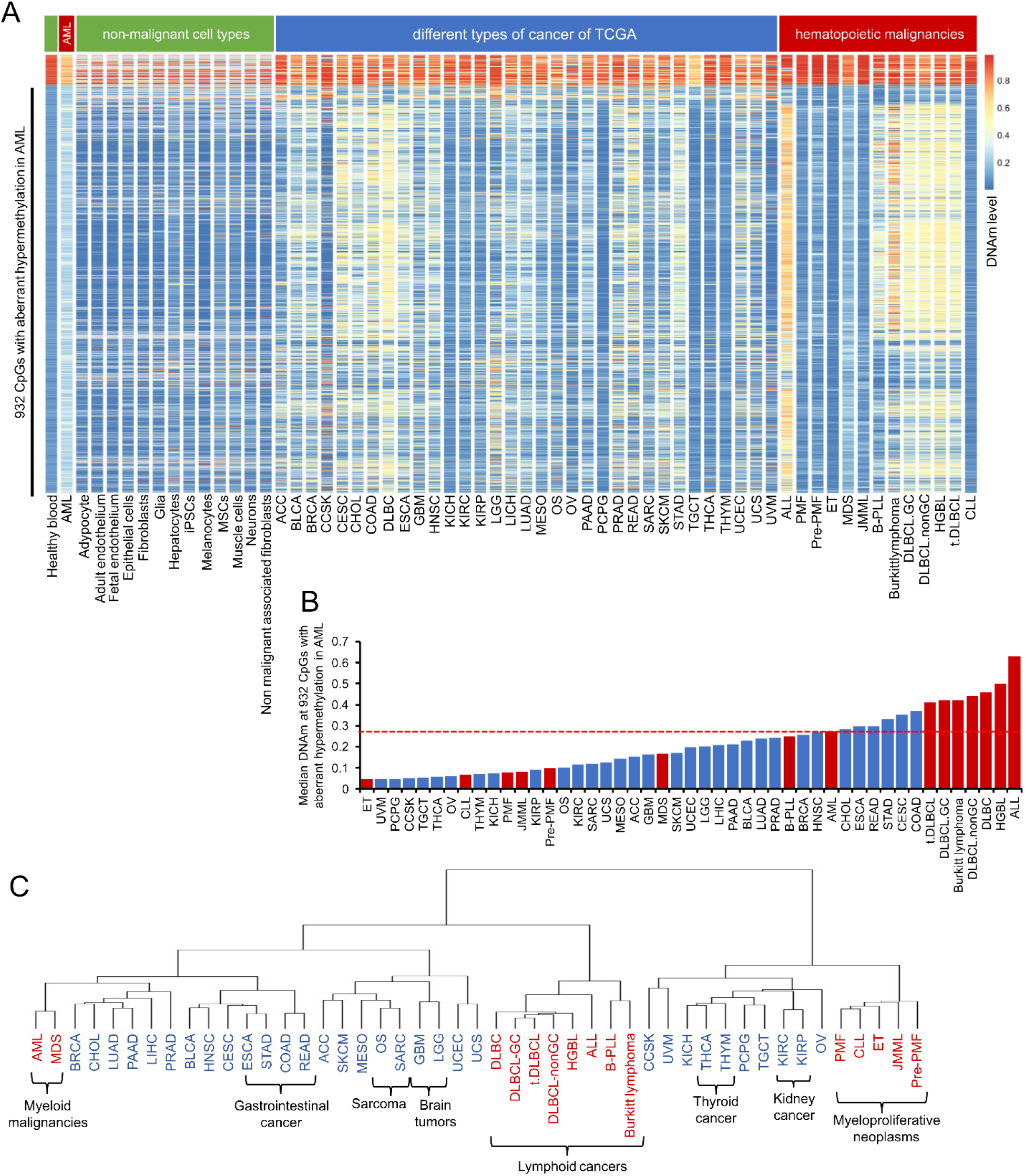
AML-associated CpGs show aberrations across different malignancies. **A)** Heatmap with mean DNAm values at the top 1000 AML associated CpGs in healthy blood samples, AML, various non-malignant cell types (green), 33 different cancer types from TCGA (blue), and 13 hemathological malignancies (red). Overall, non-malignant cell types showed low DNAm levels at the CpGs with aberrant gain of methylation in AML. In contrast, many malignancies showed a marked gain of methylation at these sites across the very different cancer types. **B)** The median DNAm level at the 932 CpGs with aberrant gain of DNAm in AML is shown across the different malignancies. Several non-hematopoietic and hematopoietic cancer types showed even more pronounced dysregulation at these sites than AML. **C)** Hierarchical clustering of the mean DNAm levels at the top 1000 AML-associated CpGs across the different malignancies. Overall, related types of malignancies clustered together.

Since the different types of cancer revealed different patterns at the 1000 AML-associated CpGs, we further analyzed if the average values within each tumor entity can be indicative for the tumor types. Hierarchical clustering of mean DNAm-values at AML-associated CpGs per disease, revealed that AML clustered most closely with myelodysplastic syndromes (MDS), while several biologically related malignancies also grouped together, suggesting that average DNAm patterns at these loci retain some degree of tissue or lineage specificity (Figure 6C). However, unlike the strong co-regulation patterns observed among individual AML patients, we did not find clearly defined co-regulated modules when using the mean DNAm levels across all patients for each of the 46 different cancer entities (Supplemental Figure 6B) – which further substantiates that the co-regulation networks can be identified between individual patients, but not between diseases.

### Epigenetic modules are preserved in non-malignant and malignant cells

Finally, we investigated whether the patient-level co-regulation network identified in AML was preserved across other malignancies. Analysis of the co-regulation matrix based on the 1000 AML-associated CpGs demonstrated highly similar co-regulated clusters in AML and MDS (Figure 7A). Comparable, although less pronounced, co-regulation patterns were also observed within the individual DNAm profiles of other cancer types, particularly in datasets with sufficient sample numbers. Importantly, when the analysis was extended to all 8,484 DNAm profiles across the pan-cancer cohort, the same higher-order co-regulated clusters remained detectable (Figure 7B). To our surprise, even in DNAm profiles of healthy blood samples (1693 samples of the training set), we observed very similar co-regulation networks as observed in the pan-cancer analysis, albeit the selected CpGs showed very little variation in absolute DNAm levels between samples (Figure 7C). Thus, also in healthy controls, there are donor-specific modulations along this co-regulation matrix – the co-regulation matrix is therefore not characteristic for malignancies, albeit extreme DNAm values are particularly found in cancer.

**Figure 7:**
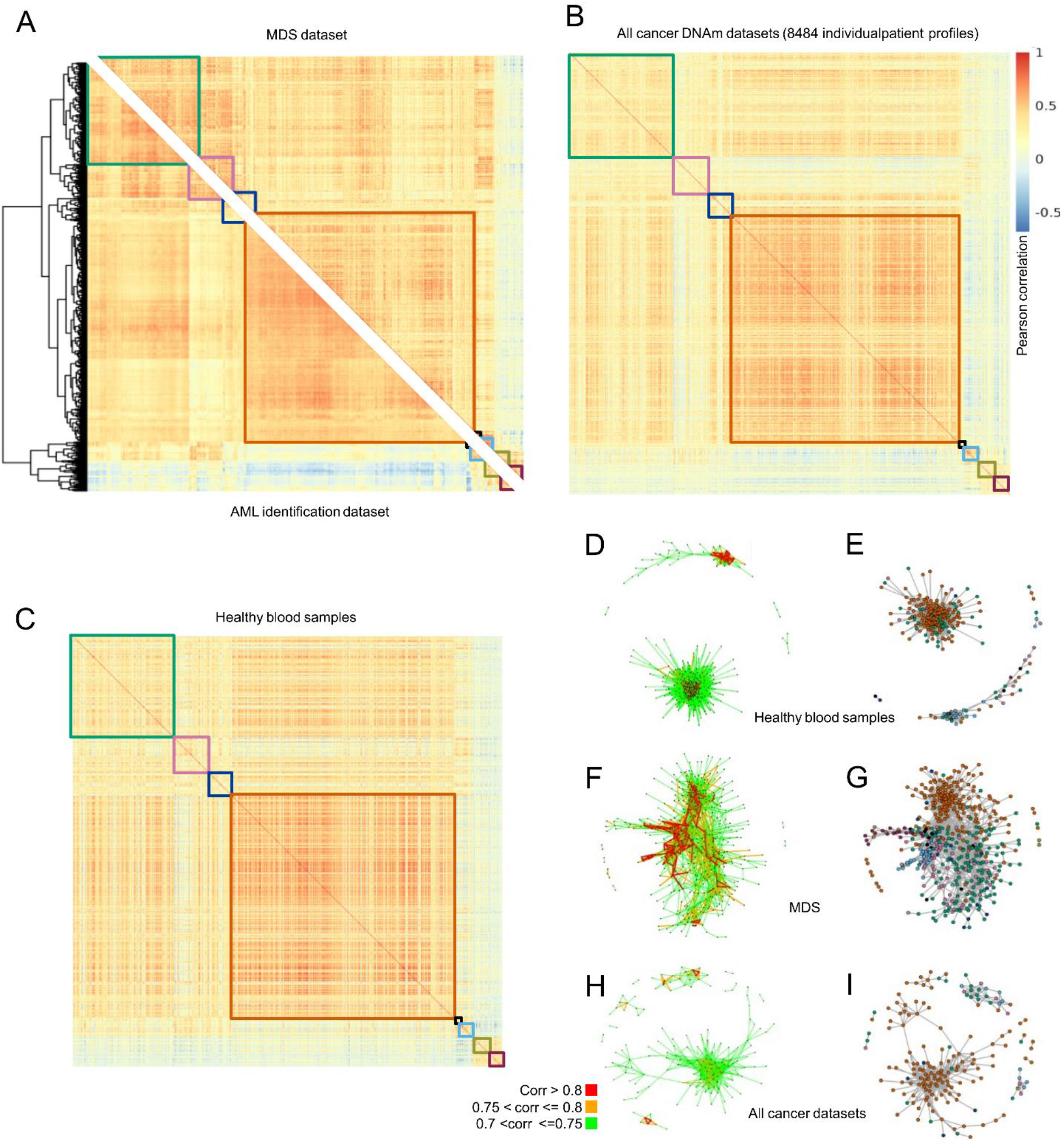
Co-regulation networks are preserved across different types of cancer. **A)** Correlation analysis (Pearson’s correlation) of DNAm at the top 1000 AML-associated CpGs within the AML identification datasets and within an MDS dataset (GSE152710; GSE221745). The correlation matrix with the eight clusters is very similar between these myeloid malignancies. **B)** When we alternatively analyzed correlation at these AML-associated CpGs in 8484 individual profiles of other hematopoietic and non-hematopoietic malignancies the correlation matrix overall revealed a very similar pattern again. **C)** Correlation matrix in 1693 normal blood samples (training set) shows very similar clusters and patterns as observed in pan-cancer data (although these CpGs were selected to have little variation in normal blood). **C-I)** Network plots for (D,E) healthy blood samples; (F,G) MDS, and (H,I) all 8484 cancers profiles. Each dot represents one of the top 1000 CpG and only pairs with a correlation value higher than 0.7 are displayed. The color coding either indicates correlation values (E,F,H), or the co-regulated clusters identified in AML (E,G,I).

We have also performed network visualization of the top 1000 CpGs for the different datasets. In healthy blood samples, the co-regulated clusters were not as distinct as in the independent AML-validation set, but showed similar features (Figure 7D,E). In MDS, we observed higher correlation coefficients between CpGs and the co-regulated clusters resembled the co-regulated clusters of AML more closely (Figure 7F,G). In the pan-cancer analysis (Figure H,I), as well as in other less-related types of cancer, the distinct modules were often conserved, but to a lesser degree than in AML. Together, these findings indicate that neither the AML-associated CpG signature nor the associated DNAm co-regulation network is restricted to AML. Rather, the recurrent co-regulation patterns across diverse malignancies point toward a shared higher-order regulatory architecture that is enhanced along co-regulatory units, which are observed also in non-malignant cells.

## Discussion

Our study places DNAm abnormalities in AML and other malignancies into a new conceptual framework: rather than representing the cumulative effect of random epigenetic drift at specific sites in the genome, these patterns reflect structured co-regulation across the genome. We describe 1) stable co-regulatory networks across independent datasets, 2) patient-specific methylation signatures that can be inferred from the epigenetic state of other chromosomes, and 3) coordinated assimilation of aberrant DNAm across homologous chromosomes. Importantly, these features are not restricted to AML, but are evident across multiple cancer types, suggesting that aberrant modulation along higher-order epigenetic networks may represent a fundamental mechanism of malignant transformation.

While many investigations have described DNAm changes in AML and other malignancies, the majority have focused on epigenetic signatures to identify biomarkers for disease classification or risk stratification without taking their potential co-regulation into account. Only a relatively small number of studies have combined gene-regulatory network analyses with epigenetic parameters to uncover functional hubs or stable states within the epigenetic landscape (29-32). Co-methylation analysis across 11 cancer types demonstrated that consistent changes in the epigenetic landscape exist in multiple cancer types (33). Furthermore, parenclitic networks of DNAm patterns have been employed for binary discrimination of cancer-positive *versus* cancer-negative samples (34). To our knowledge, a systematic co-regulatory analysis of DNAm patterns within a cancer entity, such as AML, has not yet been performed. This gap may reflect the intuitive difficulty of interpreting coherent methylation patterns without a clear mechanistic understanding of the underlying regulatory processes. From a broader perspective, however, normal development is generally characterized by remarkably tight control of the DNAm landscape, an organization that is thought to be mediated by extensive epigenetic networks (16). Disruption of these regulatory mechanisms at various levels might therefore result in aberrations that do not occur as isolated events; rather, they accumulate in a coherent, genome-wide manner.

A central aspect for studying disease associated DNAm alterations is how to choose the relevant CpG sites. Because single-cell DNAm profiling remains technically demanding, most investigations rely on bulk methylation data, and the observed epigenetic differences between AML and control samples are therefore notoriously confounded by cellular composition (8). To minimize this confounding effect, we initially filtered for CpGs that are stably methylated or stably unmethylated in healthy donors; these loci also display consistent methylation patterns across sorted normal hematopoietic subpopulations, indicating their rather cell-type-independent status. Second, we retained only those CpGs that show a large mean methylation difference between the AML cohort and the healthy controls, while allowing for substantial intra-AML variability. Our selection criteria differs fundamentally from those employed in the EVOFLUx framework that uses highly fluctuating CpGs sites in healthy controls, which function as a ‘methylation barcode’ of the tumor initiating cell (11). Furthermore, our selection strategy differs from approaches that aim to develop diagnostic biomarkers or signatures for disease stratification (18, 35, 36). In fact, we did not observe that dysregulation at our AML-associated CpG sites was clearly indicative for therapeutic outcomes, which may be partly due to increased aberrations in younger patients with less genomic aberrations.

Acute myeloid leukemia is a heterogeneous group of hematopoietic malignancies and clustering of DNAm profiles has previously been shown to stratify patients into biologically and clinically distinct subgroups according to risk, karyotype, and epigenetic driver mutations (19, 37). In contrast, our selected AML-associated CpGs did not clearly recapitulate such structured subgrouping – possibly because our selection strategy for the 1000 AML-associated CpGs captured less aspects of cellular composition or more variable CpGs. To demonstrate that the epigenetic landscape is modulated in a coherent and yet patient-specific manner, we predicted epigenetic dysregulation in a leave-one-chromosome-out analysis. These patient specific patterns demonstrate co-regulation in trans. Furthermore, we describe the assimilation of DNAm patterns on homologous alleles, which we have also previously observed in amplicon sequencing data (21). The high degree of similarity in aberrant DNAm patterns on homologous chromosomes indicates that either the patterns are copied from one allele to the other, or they are both tightly controlled by the same trans-regulating mechanisms.

Notably, we observed a surprising degree of conserved co-regulation across multiple cancer types. Predictive models trained on AML data for aberrant DNAm between independent chromosomal regions remained robust even when applied to ALL, suggesting that the co-regulation networks are not AML-specific. Previous studies have reported pan-cancer DNAm signatures (13, 38, 39), but these were typically derived from multi-cancer datasets. In contrast, our CpG selection was based exclusively on AML data, yet the resulting signatures exhibited even stronger alterations across a broad range of solid tumors. Furthermore, even in the non-malignant blood samples similar modules of co-regulation were observed, indicating that the aberrant DNAm patterns follow a similar path as in normal development – but they are clearly enhanced along the network in cancer.

The mechanisms driving aberrant DNAm in malignancies remain incompletely understood. In AML, recurrent mutations in epigenetic regulators provide a direct link between genetic lesions and DNAm patterns. For instance, mutations in *DNMT3A* are consistently associated with globally hypomethylated clusters (20, 37, 40), and we have demonstrated that aberrant hypermethylation within DNMT3A may resemble an epimutation that mimics this phenotype (41). However, a substantial proportion of AML cases – as well as many other cancer types – lack mutations in canonical epigenetic drivers. Even in mutation-positive cases, it remains unclear how such lesions could account for the complexity and coordination of genome-wide DNAm alterations in a patient-specific manner. At least, we did not observe enrichment of epigenetic driver mutations in patients with high aberrations at our AML-associated CpGs. Recent evidence suggests that somatic mutations can induce localized DNAm changes in adjacent genomic regions (15), supporting the idea that certain disease-associated DNAm patterns may arise directly from nearby genetic alterations (14). In contrast, our findings point to the existence of co-regulated epigenetic networks that appear largely independent of direct somatic mutational effects.

In analogy to gene regulatory networks (GRNs) – which are well established as key determinants of cell identity and stability (42, 43) – similar feedback-driven mechanisms may operate at the epigenetic level. Such network behavior could arise from multilayered crosstalk between regulatory processes, including different isoforms of epigenetic writers and erasers (40), methylation-sensitive binding of transcription factors that recruit DNA methyltransferases (44), interactions between DNAm and histone modifications (45), higher-order three-dimensional chromatin organization (46), regulatory contributions from non-coding RNAs such as long non-coding RNAs (lncRNAs) or PIWI-interacting RNAs (piRNAs) (47); and potentially, by assimilation during homologous recombination events (16). While the exact underlying mechanism that coordinates the cancer-associated DNAm landscape remains unknown, it becomes clear that aberrant DNAm patterns in cancer emerge from dysregulation along interconnected epigenetic networks.

Taken together, our study shows that DNAm alterations in cancer are – in contrast to genomic mutations – not the result of random events or simple epigenetic drift at isolated loci. Instead, they arise through coordinated, patient-specific genome-wide modulation through epigenetic networks, which exist also in non-malignant cells. So far, the functional relevance of these cancer-associated DNAm patterns remains unclear. However, given their genome-wide nature, such extensive epigenetic modifications are likely to have downstream functional consequences and may contribute to fundamental processes underlying malignant transformation. This may be particularly relevant in cases where no clear driver mutations or genetic abnormalities can be identified. The molecular mechanisms underlying this coordinated dysregulation of DNA methylation remain to be elucidated. However, a deeper understanding of these processes could open new therapeutic avenues. Recent studies have demonstrated that targeted epigenome editing can induce stable gene expression changes or modulate epigenetic networks, supporting its potential for future curative strategies (17, 48). If we understand the nodes that tie the epigenetic network together, it might be possible to convert a malignant into a non-malignant epigenetic phenotype.

## Methods

### Co-regulation analysis of aberrant DNAm in AML

We used DNAm profiles of peripheral blood of 737 AML patients (GSE58477, n=62; GSE62298, n=58; GSE78963, n=18; GSE80508, n=12; GSE133986, n=64; GSE159907, n=272; GSE153347, n=57, TCGA-AML, n=194), and 2390 healthy donors (GSE40279, n=656; GSE50660, n=464; GSE77716, n=573; GSE115278, n=108; GSE147740, n=1129; Supplemental Table S1) Initial quality control was performed on IDAT files using the minfi package (version 1.48.0)(49) and for subsequent preprocessing we used the sesame package (version 1.20.0): Data was normalized with noob background correction and probes with a detection p-value > 0.05 in at least one sample were removed (50, 51). Furthermore, we excluded probes associated to X or Y chromosomes, SNPs, and CpGs flagged in the b5-manifest (Illumina). Finally, we reduced our selection to CpGs that are present in the 450K and EPIC Illumina BeadChip platforms.

To select the top 1000 candidate CpGs with aberrant DNAm in AML, we initially determined the mean beta-value for healthy and AML samples, as well as the difference in mean between both groups (delta DNAm). As a first selection criterion, only CpGs with a mean DNAm level > 0.9 or < 0.1 were considered. Subsequently, we only considered those CpGs with the delta DNAm value within a 10kb window (5kb up or downstream), to exclude co-regulation with the same DMR. The remaining CpGs were then ranked according to delta DNAm values to select the top 100 or top 1000 CpGs.

Gene ontology enrichment analysis for genes associated with selected positions was performed using gometh function from missmethyl package (version 1.40.3) in R. Network visualizations were constructed using igraph package (version 2.3.1) in R, based on Pearson correlations and significance values computed with rcorr function from R package Hmisc (version 5.2). Plots were generated in R using ggplot2, ggforce and pheatmap.

### Other datasets used in the study

To estimate if AML-associated CpGs rather reflect changes in the cellular composition, we utilized DNAm profiles of purified hematopoietic subsets (GSE35069, n=48 from 6 donors) (52). To investigate if the AML-associated CpGs are also enriched at CpGs with age-associated DNAm changes, we used 656 DNAm profiles of blood samples of different aged donors (GSE40279) (24). Based on this, the Pearson’s correlation between age and DNAm was determined for each CpG, and the density distribution was subsequently plotted for different CpG subsets.

To explore the top ALL-associated CpGs, we used datasets of two studies (GSE182313, n=8; GSE147667, n=144)(25, 26). Longitudinal data from AML and ALL patients was obtained from GSE1153347 (n=33) (28) and GSE56602 (27) (n=26), respectively.

To investigate if the AML-associated epigenetic pattern was extensible to other malignancies, we interrogated in total 241 DNAm profiles of 13 hematopoietic malignancies and 8069 DNAm profiles from 33 non-hematopoietic cancer types from TCGA. Hematologic malignancies included myelodysplastic syndrome (MDS), juvenile myelomonocytic leukemia (JMML), primary myelofibrosis (PMF), prefibrotic PMF (pre-PMF), essential thrombocythemia (ET), Burkitt lymphoma, germinal center and non-germinal center diffuse large B-cell lymphoma (DLBCL.GC and DLBCL.nonGC), high-grade B-cell lymphoma (HGBL), transformed indolent B-cell lymphoma (tDLBCL), B-cell prolymphocytic leukemia (B-PLL), and chronic lymphocytic leukemia (CLL). Solid tumor entities included adrenocortical carcinoma (ACC), bladder urothelial carcinoma (BLCA), breast invasive carcinoma (BRCA), cervical squamous cell carcinoma and endocervical adenocarcinoma (CESC), cholangiocarcinoma (CHOL), colon adenocarcinoma (COAD), esophageal carcinoma (ESCA), glioblastoma multiforme (GBM), head and neck squamous cell carcinoma (HNSC), kidney chromophobe (KICH), kidney renal clear cell carcinoma (KIRC), kidney renal papillary cell carcinoma (KIRP), lower-grade glioma (LGG), liver hepatocellular carcinoma (LIHC), lung adenocarcinoma (LUAD), mesothelioma (MESO), ovarian serous cystadenocarcinoma (OV), pancreatic adenocarcinoma (PAAD), pheochromocytoma and paraganglioma (PCPG), prostate adenocarcinoma (PRAD), rectal adenocarcinoma (READ), sarcoma (SARC), skin cutaneous melanoma (SKCM), stomach adenocarcinoma (STAD), testicular germ cell tumors (TGCT), thymoma (THYM), thyroid carcinoma (THCA), uterine corpus endometrial carcinoma (UCEC), uterine carcinosarcoma (UCS), uveal melanoma (UVM), clear cell sarcoma of the kidney (CCSK), and osteosarcoma (OS). For comparison, we also utilized 346 DNAm profiles of 13 different non-malignant cell types (of 46 different studies). The information of the GSE-accession numbers and the number of samples per study are provided in Supplemental Table S3.

### Regularized multilinear models

To explore if the patient-specific patterns in aberrant DNAm could be estimated across chromosomes, we have built multiple linear regression models to predict the DNAm level of one position based on the DNAm levels of the remaining CpGs of the top 1000 selection in other chromosomes. Since the number of available samples is smaller than total parameters, the models were regularized using elastic net – and therefore not all initial parameters were finally used for the actual predictions. Models were built in R using glmnet package (v5.0), trained using one third of AML data available from Illumina 450k platform, and validated on the rest of AML data: the remaining two thirds of data from Illumina 450k platform and all available data from EPIC platform. All plots were generated in R using ggplot2 (v 4.0.0) and ggforce (v0.5.0).

### Generation and analysis of long-read nanopore sequencing data

Peripheral blood samples from 10 AML patients and 10 healthy controls were obtained from RWTH central biomaterial bank (cBMB) after informed and written consent from donors (approval ID: EK206_09). DNA was extracted using DNA Blood Micro-Kit Qiamp (Qiagen) and sequenced on a Oxford Nanopore Technologies PromethION P24 sequencer. To achieve higher coverage at our sites of interest, we used adaptative sampling for the 10kb window around the top 100 AML-associated CpGs. Basecalling, demultiplexing, and alignment (hg38 annotation) were performed using dorado (v0.7.2). Methylation calls were extracted using Modkit (v0.4.0 for healthy samples and v0.4.2 for AML samples). One control sample was not considered for further analysis due to bad quality.

## Additional information

### Data availability

We have utilized publically available DNAm profiles of many different studies, as indicated in the text (e.g. Supplemental Table S1 and S3, with 3667 and 8830 DNAm profiles, respectively). The adaptive nanopore sequencing data are available upon request and approval of the current data protection regulation at RWTH Aachen medical school.

### Author Contributions

M.V.B., S.L., T.S. and W.W. contributed to the experimental design. M.V.B., S.L., F.K., C.-C.K. and J.-F.P.-C. performed the bioinformatic analysis. M.V.B. and T.S. performed network analysis. M.V.B. and F.K. carried out the nanopore sequencing experiments. E.J. contributed viable materials. W.W. initiated and supervised the study and wrote the initial draft of the manuscript. All authors revised and approved the final version of the manuscript.

### Conflicts of Interest

W.W. is cofounder of the company Cygenia GmbH (www.cygenia.com) that provides services for epigenetic analyses to other scientists. Apart from this the authors have no competing interests to declare.

### Funding

This study was particularly supported by the Else Kröner-Fresenius Stiftung (2025_EKSE.35 to W.W.) and by the Deutsche Forschungsgemeinschaft (DFG; WA1706/17-1/561150360 to W.W. and T.S.). Furthermore, we acknowledge financial support by DFG (WA1706/12-2/417911533; WA1706-14-1/458369830; SFB 1506-1/450627322; and GRK 2415/2/363055819, all to W.W.), the Federal Ministry of Education and Research (VIP+: 03VP11580, to W.W.), and by the José Carreras Foundation (DJCLS 03 R/2024 to W.W.).

## Acknowledgement

We acknowledge the Biobank of RWTH Aachen University Medical School for colleting the AML samples, and the computing resources granted by RWTH Aachen University. We thank Sebastian Giesselmann for his support with the preparation of Nanopore sequencing libraries.

