## Supplemental figures S1-S6, and tables S1 and S3 for "Aberrant DNA methylation is co-regulated across the genome in leukemia and other types of cancer"

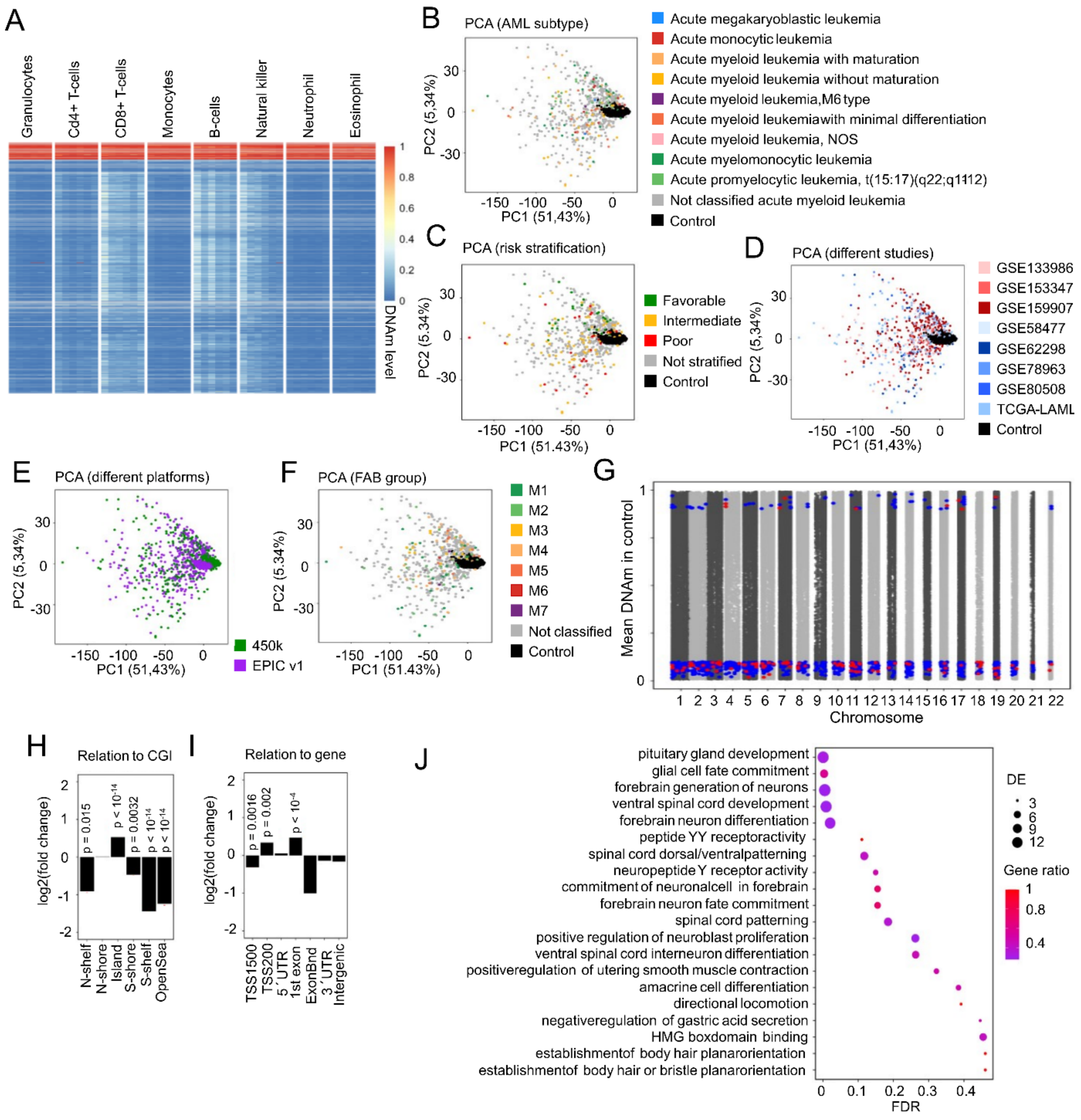


**Supplemental Figure S1. Further analysis of AML-associated CpGs.**

**A)** Heatmap of DNAm levels at the top 1000 AML-associated CpGs in different sorted hematopoietic subsets (GSE35069). **B-F)** Principal component analysis based on the beta-value of the 1000 AML-associated CpGs in datasets of AML and healthy controls. While the control samples clustered closely together, we did not see a clear clustering according to (B) AML subtype, (C) risk stratification of AML, (D) the different studies that the DNAm profiles were derived from, (E) the different Illumina BeadChip platforms (450k or EPICv1), or (F) french-american-british classification (FAB) subtype (samples with missing information are indicated in grey). **G)** Manhattan plot displaying the distribution of top 100 (red) and top 1000 (blue) AML-associated CpGs across the different autosomal chromosomes. **H,I)** Enrichment of the top 1000 AML-associated CpGs in relation to (H) CpG islands (CGI), and (I) in relation to genes. The log2 fold change was determined as compared to all CpGs on the microarray and P-values for enrichment are indicated (determined by Fisher exact test with Bonferroni correction for multiple testing). **J)** Gene ontology classification of genes associated with the top 1000 AML-associated CpGs. From the significantly enriched terms (adjusted p-value < 0.05), the 20 terms with the highest ratio are depicted.

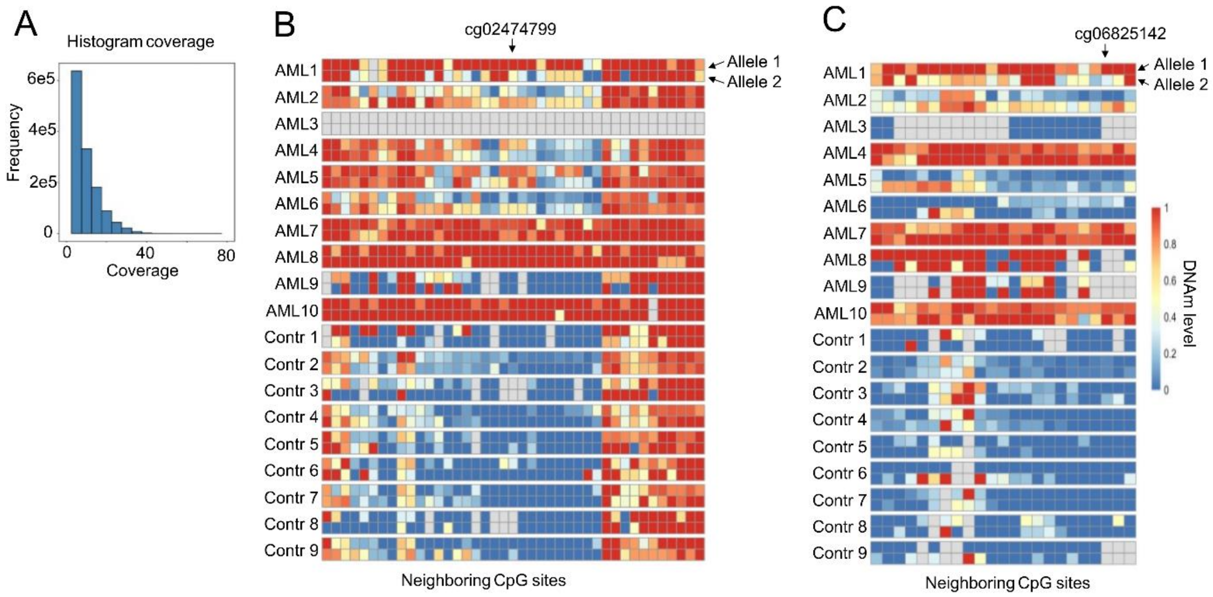


**Supplemental Figure S2: Nanopore sequencing analysis aberrant DNAm in AML.**

**A)** Adaptive nanopore sequencing was performed at the top 100 AML-associated CpGs and the histogram displays the coverage within individual samples (9 healthy control samples and 10 AML samples) at the 10kb genomic region surrounding these CpGs. **B,C)** The DNAm patterns on both alleles were analyzed in phased nanopore sequencing data and exemplarily depicted for (B) cg02474799, and (C) cg06825142. The relevant CpG site is indicated and DNAm levels at all neighboring CpGs within a 4kb window are demonstrated. Each row corresponds to the different alleles of the corresponding samples.

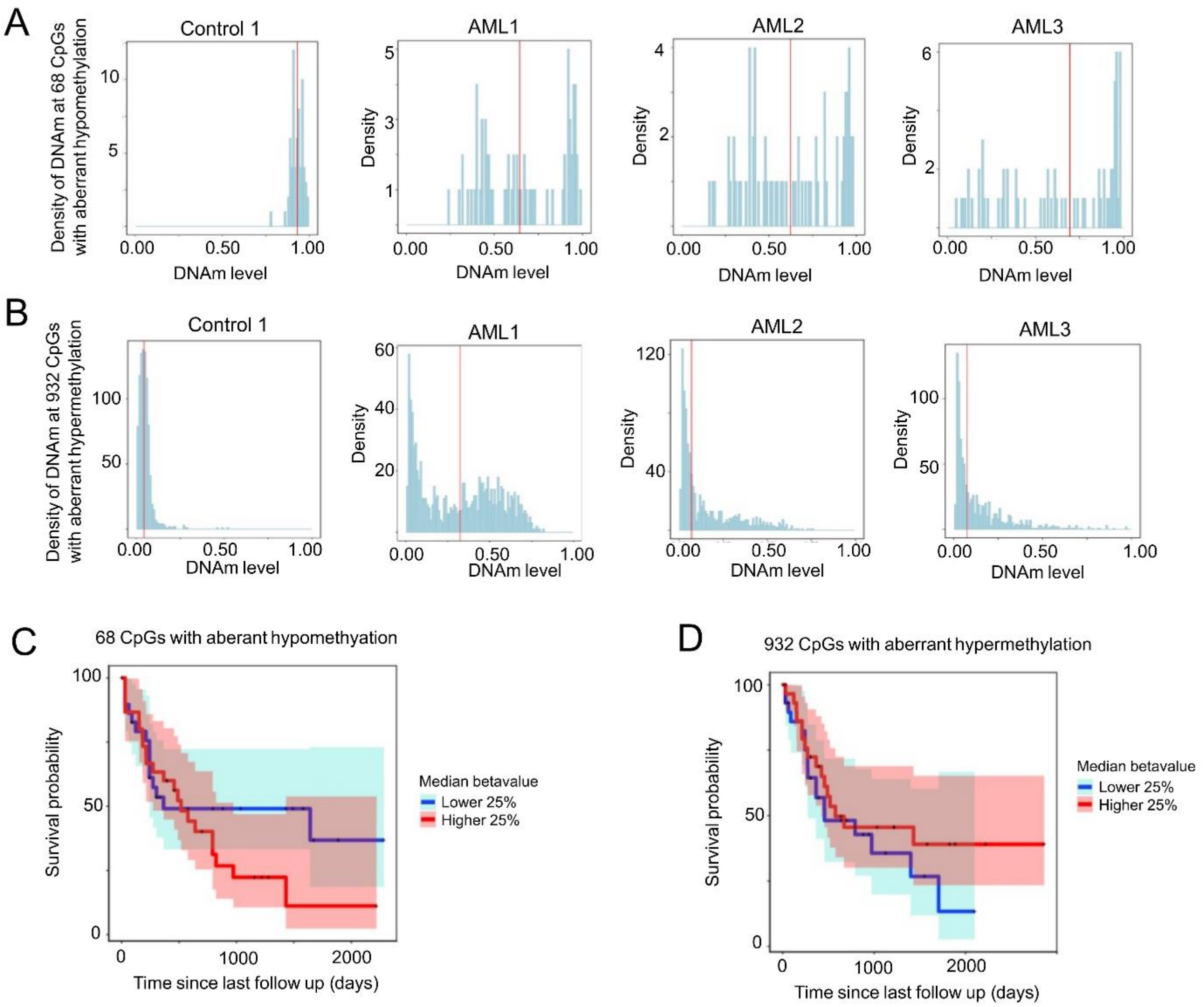


**Supplemental Figure S3: Heterogeneity of DNAm levels at AML-associated CpGs.**
**A,B)** Density plot of DNAm levels (beta-values) for an exemplary healthy sample (left) and three exemplary AML samples, at (A) the 68 CpGs with aberrant loss of DNAm, or (B) the 932 CpGs with aberrant gain of DNAm. These plots depict the heterogeneity of DNAm levels at these AML-associated CpGs within each donor. The median of these DNAm levels is indicated as red line, and this was used as a surrogate for the overall dysregulation within each sample. We have alternatively investigated association of patient specific parameters with the top 10% of CpGs or the mean values, and the results were very similar (not demonstrated). **C,D)** Kaplan-Meier plots showing the survival probability in the TCGA-AML cohort. The quartiles with the lowest (blue) or highest (red) median DNAm at either the CpGs with aberrant loss of DNAm (C), or aberrant gain of DNAm in AML (D) are compared. In tendency, both curves indicate that samples with more epigenetic dysregulation have a better prognosis, but the analysis did not reach statistical significance.

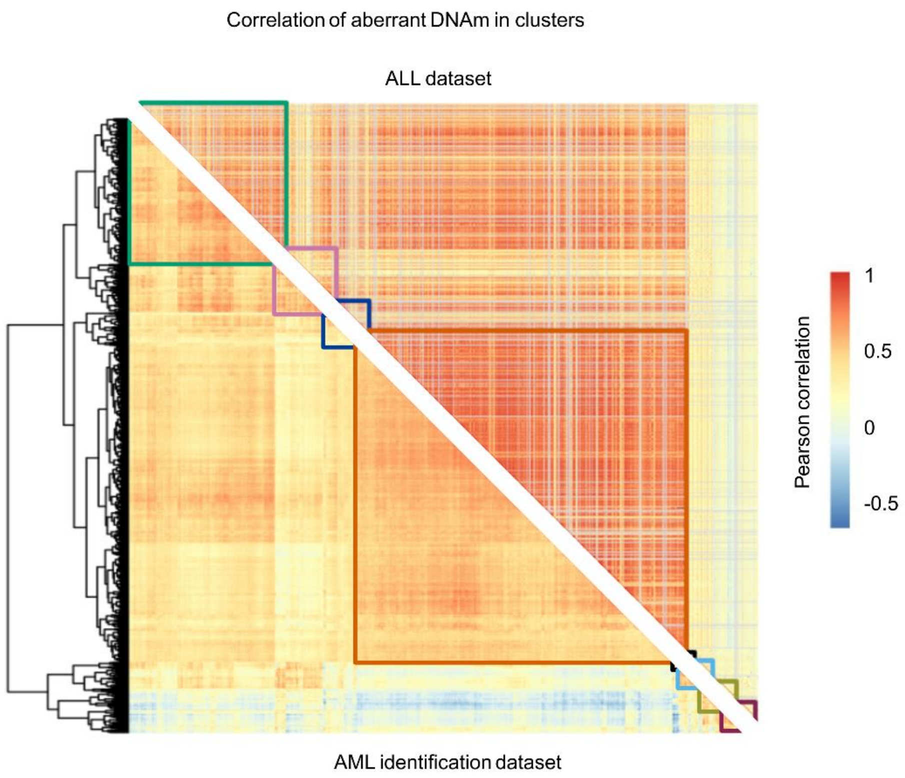


**Supplemental Figure S4: Co-regulation network at AML-associated CpGs within ALL samples.**
**A)** Correlation analysis (Pearson´s correlation) of DNAm at the top 1000 AML-associated CpGs within the AML identification datasets and within ALL datasets (GSE182313 and GSE147667). Several pairs of CpGs show even higher correlation across the ALL patients as compared to AML patients and the eight clusters are overall preserved even across the very different types of leukemia.

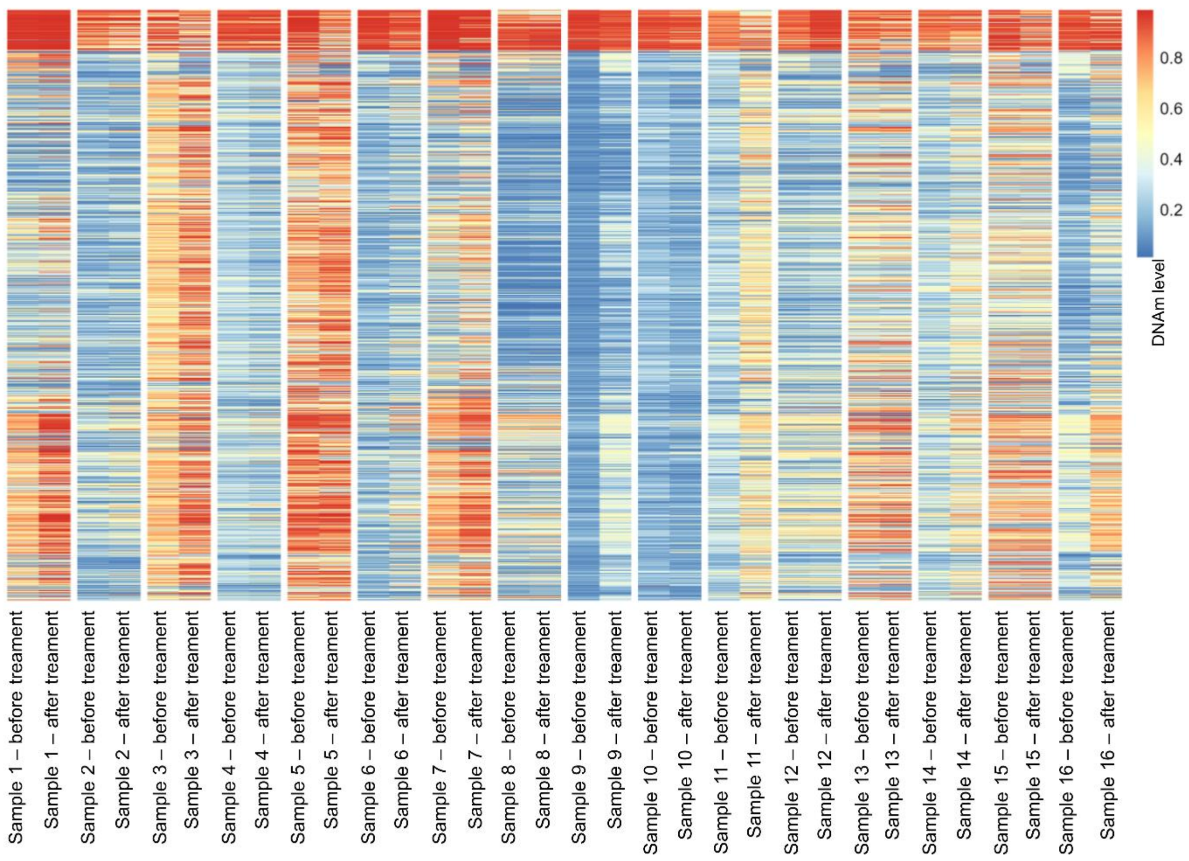


**Supplemental Figure S5: Aberrant DNAm patterns in AML increase during disease.**
Heatmap of the DNAm levels at the top 1000 AML-associated CpGs (in the same order as for heatmap in Figure1D) for 16 non-responders before treatment and after treatment (GSE1153347). This comparison demonstrates that the aberrant patient-specific DNAm patterns become even more prominent during disease. In responders this effect was not evident, since the fraction of malignant cells decreases.

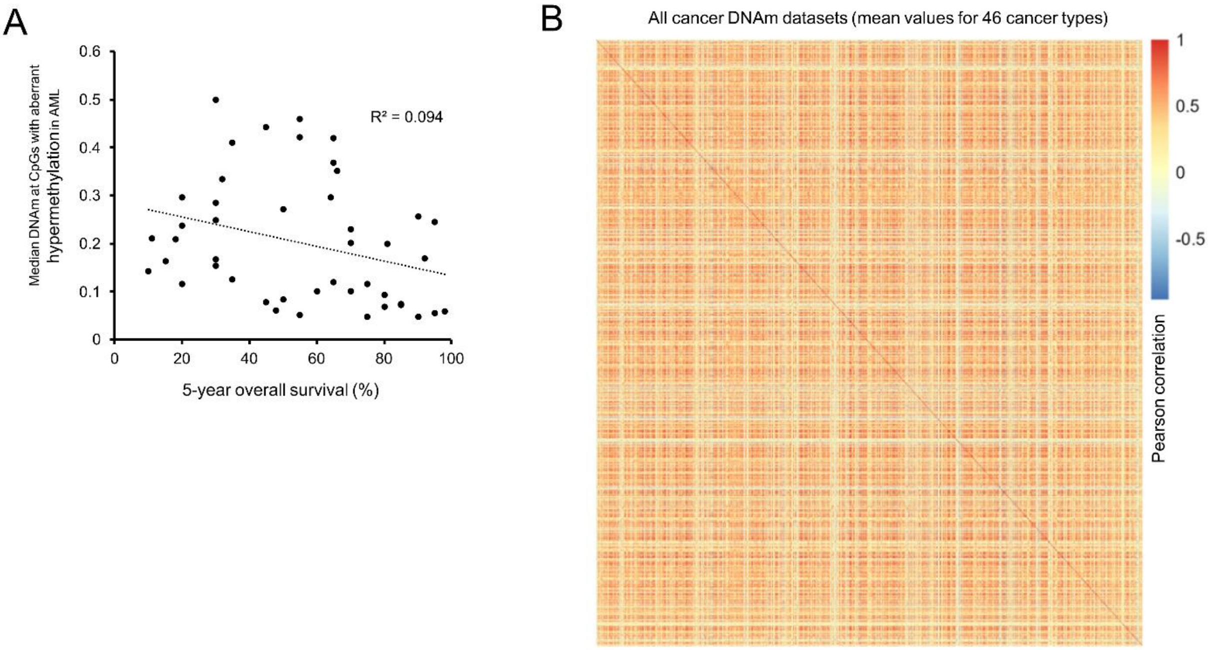


**Supplemental Figure S6: Aberrant DNAm patterns across different types of cancer.**

**A)** Regression between the general 5-year survival rate for individual cancer types and the median DNAm level at the 932 CpGs with aberrant gain of DNAm in AML. Each dot represents one proliferative disease (in total 57 cancer types). The 5-year survival rates are based on a literature search that was supported unbiased by ChatGPT. In tendency, the more aggressive cancer types, with a lower percentage of 5-year OS, showed more aberrant gain of DNAm at these sites. **B)** The mean DNAm values were calculated within each of the 57 cancer types for the top 1000 AML-associated CpGs (also depicted in Figure 6A). The correlation heatmap between pairs of CpGs (same order as in heatmap from Figure 1E) did not reveal the clusters that are observed when analyzing the individual patients. This demonstrates that the co-regulation networks are only reflected based on the heterogenous patient-specific patterns, but not based on the mean values of the different diseases (because the co-regulation networks are averaged across all patients within each cancer entity).

### Supplemental tables

**Supplemental Table S1: Public datasets of AML and healthy blood methylation array profiles used for this study**

| GEO Accession Number | Number of samples | Sample type | Illumina platform | Assignment to datasets |
|---|---|---|---|---|
| GSE58477 | 62 | AML | 450K | Selection |
| TCGA-LAML | 194 | AML | 450k | Selection |
| GSE78963 | 18 | AML | 450k | Selection |
| GSE80508 | 12 | AML | 450k | Selection |
| GSE62298 | 58 | AML | 450k | Selection |
| GSE40279 | 656 | Healthy | 450K | Selection |
| GSE50660 | 464 | Healthy | 450K | Selection |
| GSE77716 | 573 | Healthy | 450K | Selection |
| GSE133986 | 64 | AML | EPIC v1 | Validation |
| GSE159907 | 272 | AML | EPIC v1 | Validation |
| GSE153347 | 57 | AML | EPIC v1 | Validation |
| GSE115278 | 108 | Healthy | EPIC v1 | Validation |
| GSE147740 | 1129 | Healthy | EPIC v1 | Validation |

**Supplemental Table S2. Top 1000 AML-associated CpGs**

This table is provided as a separate EXCEL file. It provides CpG ID; associated genes, mean DNAm in healthy, meanDNAm in AML, and mean difference.

**Supplemental Table S3: Additional datasets of non-malignant cells, hematopoietic malignancies, and solid cancer**

| Name | ShortCut | GEO/TCGA Accession Number | Number of samples |
|---|---|---|---|
| Adipocytes | | GSE58622; GSE122126 | 15 |
| Endothelial cells (adult) | | GSE122126; GSE74877; GSE87177; GSE34486 | 19 |
| Endothelial cells (fetus) | | GSE103253; GSE106099; GSE40699; GSE82234; GSE99716 | 42 |
| Epithelial cells | | GSE109042; GSE122126; GSE40699; GSE74877; GSE85566 | 31 |
| Fibroblasts | | GSE107226; GSE111396; GSE40699; GSE41933; GSE52025; GSE65078; GSE68134; GSE68851; | 73 |

| | | | |
|---|---|---|---|
| | | GSE74877; GSE77135; GSE86258; GSE95096 | |
| Glia | | GSE74486; GSE79144 | 17 |
| Hepatocyte | | GSE122126; GSE40699; GSE60753 | 17 |
| Induced pluripotent stem cells | iPSCs | GSE51921; GSE59091; GSE65078; GSE68134 | 25 |
| Melanocytes | | GSE74877 | 3 |
| Mesenchymal stroma cells | MSCs | GSE41933; GSE52112; GSE79695; GSE87797 | 67 |
| Muscle cells | | GSE53302 | 6 |
| Neuron | | GSE122126; GSE74486; GSE79144; GSE98203 | 28 |
| Nonmalignant tissue associated fibroblast | | GSE86829 | 3 |
| Adrenocortical carcinoma | ACC | TCGA | 80 |
| Clear cell sarcoma of the kidney | CCSK | TCGA | 11 |
| Osteosarcoma | OS | TCGA | 86 |
| Bladder urothelial carcinoma | BLCA | TCGA | 418 |
| Breast invasive carcinoma | BRCA | TCGA | 793 |
| Cervical squamous cell carcinoma and endocervical adenocarcinoma | CESC | TCGA | 307 |
| Cholangiocarcinoma | CHOL | TCGA | 36 |
| Colon adenocarcinoma | COAD | TCGA | 312 |
| Lymphoid neoplasm diffuse large B-cell lymphoma | DLBC | TCGA | 48 |
| Esophageal carcinoma | ESCA | TCGA | 185 |
| Glioblastoma multiforme | GBM | TCGA | 140 |
| Head and neck squamous cell carcinoma | HNSC | TCGA | 473 |
| Kidney chromophobe | KICH | TCGA | 66 |
| Kidney renal clear cell carcinoma | KIRC | TCGA | 324 |
| Kidney renal papillary cell carcinoma | KIRP | TCGA | 275 |
| Brain lower grade glioma | LGG | TCGA | 516 |
| Liver hepatocellular carcinoma | LIHC | TCGA | 377 |
| Lung adenocarcinoma | LUAD | TCGA | 473 |
| Mesothelioma | MESO | TCGA | 87 |
| Ovarian serous cystadenocarcinoma | OV | TCGA | 10 |
| Pancreatic adenocarcinoma | PAAD | TCGA | 184 |
| Pheochromocytoma and paraganglioma | PCPG | TCGA | 179 |
| Prostate adenocarcinoma | PRAD | TCGA | 502 |
| Rectum adenocarcinoma | READ | TCGA | 98 |
| Sarcoma | SARC | TCGA | 261 |
| Testicular germ cell tumors | TGCT | TCGA | 150 |
| Thyroid carcinoma | THCA | TCGA | 507 |
| Thymoma | THYM | TCGA | 124 |

| | | | |
|---|---|---|---|
| Uterine corpus endometrial carcinoma | UCEC | TCGA | 438 |
| Uterine carcinosarcoma | UCS | TCGA | 57 |
| Uveal melanoma | UVM | TCGA | 80 |
| Skin cutaneous melanoma | SKCM | TCGA | 104 |
| Stomach adenocarcinoma | STAD | TCGA | 395 |
| B-cell promyelocytic leukemia | B-PLL | GSE279602 | 20 |
| Burkit lymphoma | | GSE269421 | 6 |
| Chronic lymphocytic leukemia | CLL | GSE206842 | 6 |
| Germinal center diffuse large B-cell lymphoma | DLBCL.GC | GSE255869 | 19 |
| Non germinal center diffuse large B-cell lymphoma | DLBCL. nonGC | GSE255869 | 17 |
| Essential thrombocythemia | ET | GSE277841 | 2 |
| High grade B-cell lymphoma | HGBL | GSE255869 | 6 |
| Juvenile myelomonocyte leukemia | JMML | GSE237299 | 41 |
| Myelodysblastic syndrome | MDS | GSE152710; GSE221745 | 78 |
| Primary myelofibrosis | PMF | GSE277841 | 27 |
| Prefibrotic primary myelofibrosis | pre-PMF | GSE277841 | 1 |
| Transformation of indolent B-cell lymphoma | tDLBCL | GSE255869 | 13 |
| Acute lymphoid leukemia | ALL | GSE182313; GSE147667 | 152 |

The color code indicates non-maligant cell types (green), solid tumors from TCGA (blue), and hematopoietic malignancies (red; in addition to AML)
